# BioBankRead: Data pre-processing in Python for UKBiobank clinical data

**DOI:** 10.1101/569715

**Authors:** D. Schneider-Luftman, W. R. Crum

## Abstract

**Motivation:** UKBiobank collects health-related data from 500,000 volunteers and is widely used by medical researchers. However, the data is supplied in a custom compressed format and its preparation for analysis is cumbersome and time-consuming. This Python package automates the extraction of selected UKBiobank data, for easy integration in an analysis pipeline.

**Features:** The functions provided within this Python package reduce the number of steps, as well as human and computational time, required for extraction and preparation of the data for analysis. It is executable through command line, is easily installed on any platform and requires no prior knowledge of Python.

**Application:** BiobankRead is used for an analysis of dietary lifestyles and cardio-vascular outcomes. A large range of dietary, phenotypical, lifestyle and vascular outcomes is extracted and pre-processed. Significant associations are found between non-meat-eating and lower blood pressure / reduced risk of hypertension.

**Availability:** The Python package BiobankRead is freely available under the GNU General Public License (version 3 or later). It can be downloaded from GitHub (https://github.com/saphir746/BiobankRead-Bash), where example scripts and detailed instructions are also available.

## Introduction

UKBiobank (http://www.ukbiobank.ac.uk/) is an important source of population-scale health data of interest to researchers across medicine. It gathers an extensive range of clinical, physiological, genomic and imaging variables from over 500,000 subjects. However, extracting and pre-processing this data is complex, time-consuming and susceptible to human error. Bespoke software for analysing UK Biobank data is starting to appear, most notably PHESANT [Millard L.A.C. et al., 2017]. Similarly, there is a growing interest in software to decode, extract, preprocess and combine sets of phenotype variables. To our knoweldge, only ukbtools [Hanscombe K. B., et al, 2017] provides some functionality for extraction and preparation of data frames for UKBiobank clinical and genomic data, for use in PLINK.

BiobankRead is a Python package designed to streamline the pre-processing of UK Biobank data, returning readily usable data-frames for any subset of variables. It provides fast pre-processing tools for clinical and phenotypical data. BiobankRead currently supports main project and HES records data (Hospital Episode Statistics, http://www.hscic.gov.uk/hes) but does not provide tools for genomic or imaging data.

In order to demonstrate the functionality of the package, its use is illustrated on an analysis of dietary lifestyles variables and cardio-vascular outcomes. In total, over 60 variables – across phenotypic, lifestyle, health outcomes and food intake fields – were extracted and pre-processed in order to facilitate the analysis [appendix 1]. Dietary lifestyles have seldom been researched in UKBiobank [Bradbury et al. 2017, Tong et al. 2018], which may be explained by inherent complexity of dietary and food intake data.

## The data

### UKBiobank data

UK Biobank recruited 500,000 people aged between 40-69 years in 2006-2010 from across the UK. The data is available, upon application, to researchers conducting health-related research for public benefit [Sudlow, C. et al., 2015; Ollier, W., Sprosen, T. & Peakman, T., 2005].

Approved investigators have access to data downloaded in an encrypted compressed .enc format as part of a defined project. Unpacking (http://biobank.ctsu.ox.ac.uk/crystal/docs/UsingUKBData.pdf) generates two files, a .csv (comma-separated-variable) file and a .html file. The .csv file contains the project data while the .html file explains how the .csv file is structured. Conventionally, researchers can open and read the .html file, look for a variable, find its corresponding column number in the .csv file, then extract that column number using STATA, R, Python or other tools. For each variable, there are between 1 and 28 measurements, across 3 assessments (baseline, first and second re-visits). Hence, for each variable, there can be up to 84 associated columns. In most analysis using this data, a large number of variables will be required, and it is essential that their extraction and pre-processing is as seamless and accurate as possible. There is evidently a need for an automated process.

### HES records

HES (Hospital Episode Statistics) data records incidents of hospitalisation in England from 1987. It uses the World Health Organization’s ICD (International Classification of Diseases) and the OPCS (Office of Population, Censuses and Surveys). It contains information about dates of admission/release/operation, diagnosis codes, in ICD10 (or ICD9 if prior to 2000), as well as operations & procedure (OPCS) codes when applicable. The aim of researchers using HES data is to find instances of specific diseases or incidents and link them to the subject’s lifestyle, genomic profile or imaging results (ex: Cullen, B. et al., 2017; Celis-Morales, C.A. et al., 2017). To achieve this, it is essential to devise fast and efficient algorithms that can retrieve results for any condition across all the coding protocols.

## Implementation

The package is written in Python, which is widely used in biomedical computational pipelines and runs on most common computing platforms. BiobankRead provides functionality for loading, sanitizing, preprocessing and extracting phenotype data from UK Biobank using Pandas [McKinney, W., 2010] data-frames and Beautiful Soup 4 (BS4) [Richardson, 2012] for data-parsing. BiobankRead defines a class with associated methods for searching, extracting and manipulating phenotype variables, including averaging over visits, removal of confounds and outliers, correlation and covariance calculation and output in universally readable Comma-Separated-Value (CSV) files. Methods are also available for processing associated Hospital Episode Statistics (HES), ICD10 codes and self-reported illness data.

We provide high-level scripts which hide the details of the class interface and, for most users, provide all the functionality they need: (i) ***extract_variables*** (extract specific variables with conditional filtering), (ii) ***extract_death*** (output cases with specific mortality causes), (iii) ***extract_SR*** (output cases with specific Self-Reported conditions), (iv) ***extract_HES*** (output hospital admission episode dates for cases with specified conditions). All scripts require the standard Biobank data input files in .csv and .html. We provide an additional script **search_var** to identify variables of interest with keyword combinations.

### Other operations

BiobankRead features other processing options which can be called directly from Python or used as options in the high-level scripts. **extract_variables** includes options for retrieving first visit data only, averaging over visits, removing cases with missing data and removing outliers. **extract_death** includes options for considering primary and/or secondary causes of death. **extract_SR** includes options for cancer / non-cancer and whether to examine first visit data only. **extract_HES** includes options for retrieving first visit data only, outputting episode start or admission date and marking hospital visits before and after baseline Biobank assessment. BiobankRead also extracts and understands data-codings for categorical variables (example: YES / NO / OCCASIONALLY categorical answers about cigarette smoking).

### Custom variable design

One of the most significant appeals of UKBiobank is its extensive wealth of data. A common practice is to formulate new variables using available ones, in order to investigate complex traits. For instance, in [Guo, W. et al., 2015], a measure of physical activity in women pre and post menopause is derived using several physical activity variables, and is expressed in MET-h/week. All these steps can easily be performed using BiobankRead. More complicated manipulations of similar nature are often required in epidemiology studies, notably when working with food intake data. This is illustrated in the “Usage” section below.

## Usage – Diet choices and blood pressure

In this paper we demonstrate the usage of BiobankRead on an example involving dietary choices and vascular health.

Nutrition can have a substantial impact on cardiovascular health. In particular, a clear link has been established between red meat / processed meat intake and colorectal cancer [Aykan, 2015], whereas increased intake of vegetables, particularly cruciferous vegetables (broccoli, cauliflower, cabbage…), have protective effects against cardiovascular disease [Zhang et al., 2011]. In light of this, and based on concerns for environmental protection / animal welfare, some dietary lifestyles have gained substantial popularity, notably vegetarianism and veganism. In the UK, it is estimated that vegetarianism expanded by 40% and veganism by 261.33% between 2006 and 2016. It is hypothesized that these dietary choices offer a range of health benefits, notably for vascular health, however this has been little studied in the literature [Crowe et al, 2013]. To our knowledge, this has only been examined in UKBiobank across physiological characteristics and protein intake [Bradbury et al. 2017, Tong et al. 2018]. We explore here the intersection between these dietary variables and vascular outcome data, after extracting all relevant data fields with BiobankRead.

Four dietary groups are defined based on their non-intake of various foods: omnivorous (476,362 - 95.7%), pescatarian (11,381 - 2.3%), vegetarian (9,464 – 1.9%) or vegan (578 – 0.1%). Outcomes of interest are: systolic blood pressure (SBP) and hypertension status (hBP). For this analysis, hBP is defined for any subject with at least one of the following: SBP>140, DBP>90, taking medication for hypertension, or high blood pressure diagnosed by doctor. The outcomes of interest are analyzed against the dietary lifestyle groups and a range of confounders in a linear regression (SBP) and logistic regression (hBP):

- Model 1: outcome ∼ Diet group + Sex + Birthyear
- Model 2: Model 1 + BMI + Sedentary + Alcohol intake + current smoking status

All statistical analyses were performed in R v. 3.4.4.

Results are displayed in table 2. Results are expressed as regression coefficients, std. Deviation and p-value for the LM model on Systolic blood pressure, with reference coefficient set to 0 for the omnivorous group. Results are expressed in odds ratio (ORs), confidence intervals and p-value for the logistic model on hypertension, relative to the omnivorous group (OR=1). There is a significant reduction in systolic blood pressure across all non-meat-eating groups (consistent with findings of [Tong et al., 2018]), as well as a significantly reduced odds for hypertension. The sharpest decrease in SBP and hypertension incidence is consistently observed amongst the vegan group, though the significance of results for this group is moderate due to its small sample size.

**Table 1:** descriptive summary of BiobankRead functions. All input are passed as command line options, and output directly returned to a directory specified by the user

| SCRIPT | BASIC USAGE (other options are available) | BASIC EXAMPLE |
| --- | --- | --- |
| **search_var** | --keywords key1 ... keyN<br>--match and / or | --keywords age blood<br>--match and |
| **extract_variables** | --vars variable1 ... variableN<br>--filter condition1 ... conditionM | --vars "Birth weight"<br>--filter "Birth weight>3" |
| **extract_death** | --codes code1 ... codeN | --codes C34 C42 |
| **extract_SR** | --disease disease_name | --disease "lung cancer" |
| **extract_HES** | --tsv HES_data_file<br>--codes code1 ... codeN | --tsv ukb.tsv<br>--codes C49 |
| **ALL** | --out output_path | --out results.csv |

**Table 2:** regression analysis results for diet groups against Systolic Blood Pressure (SBP) (top) and hypertension (HBP) status (bottom). Model 1: Diet group + Birthyear + Sex; Model 2: Model 1 + BMI + sedentary^*^ + Alcohol intake^2^ + Current Smoking status. Omnivorous group is used as reference for all models and outcomes, coefficients and ORs are expressed in terms of relative difference / “risk” to the omnivorous group.

|  | Pescatarian | Vegetarian | Vegan |
| --- | --- | --- | --- |
| ***SBP Lin. Regr.*** |  | **Model 1** |  |
| Coeff. | -2.9 | -1.85 | -3.9 |
| Std. dev | 0.17 | 0.19 | 0.74 |
| P. value | <2.1e-16 | <2.1e-16 | 1.2e-07 |
|  |  | **Model 2** |  |
| Coeff. | -1.66 | -0.56 | -1.7 |
| Std. dev | 0.17 | 0.19 | 0.74 |
| P. value | <2.1e-16 | 2.8e-03 | 0.02 |
| ***HBP Logistic*** |  | **Model 1** |  |
| OR | 0.67 | 0.8 | 0.6 |
| C. I | [0.64, 0.7] | [0.76, 0.83] | [0.5, 0.71] |
| P. value | <2.1e-16 | <2.1e-16 | 2.5e-08 |
|  |  | **Model 2** |  |
| OR | 0.83 | *0.96* | 0.78 |
| C. I | [0.8, 0.87] | *[0.92, 1.01]* | [0.65, 0.95] |
| P. value | 3.6e-16 | *0.43* | 0.012 |

*: hrs / day spent doing “sedentary” activities: watching TV, sat at desk and/or driving

2: categories of alcohol intake: 1-everyday, 2 – most days, 3 - once/twice a week, 4 - once/twice a month, 5 – occasional, 6 - never

## Discussion

BiobankRead was designed as a toolset to ease and accelerate the pre-processing of epidemiology data in UKBiobank, and is particularly relevant for devising complex, custom-made variables. It can extract variables of interest in a fast and automated way, requiring only minimal command line inputs from the users. Its strength also lies in its functions for HES data manipulation.

BiobankRead is designed for researchers who need to repeatedly manipulate and customise UKBiobank data, notably in conjunction with Hospital Episode Statistics. BiobankRead is designed to handle phenotypical, clinical and HES data, and does not currently offer functionalities for imaging and genomic data. BiobankRead offers functionalities that complement those of PHESANT and ukbtools. The PHESANT package performs automated pre-processing and associated phenome analysis., while ukbtools is specifically designed in R to pre-process data GWA. BiobankRead is a lower-level tool which allows more customisation of confounding variables, pre-processing of HES, Self-Reported and mortality data. An important distinction, showcased above, is the ability of BiobankRead to synthesis complex variables and health outcomes.

UKBiobank is a resource of growing and considerable importance to the bio-medical research community, and the ability to handle this data set effectively and succinctly is essential for the discovery of new associations between the genome, lifestyle factors, phenotypes and clinical outcome, but also for reproducibility. BiobankRead allows for the transformation of UKBiobank data into fully-customisable data frames, optimized for analyses.

## Acknowledgement

This research has been conducted using the UK Biobank Resource, via applications number 236 and 10035. Many thanks to Dr A. Berlanga for his valuable contributions to the code.

## References

Sudlow, C. et al., 2015. UK Biobank: An Open Access Resource for Identifying the Causes of a Wide Range of Complex Diseases of Middle and Old Age. PLoS Medicine, 12(3).

Ollier, W., Sprosen, T. & Peakman, T., 2005. UK Biobank: from concept to reality. Pharmacogenomics, 6(6), pp.639–646.

Cullen, B. et al., 2017. The “cognitive footprint” of psychiatric and neurological conditions: cross-sectional study in the UK Biobank cohort. Acta Psychiatrica Scandinavica, 135(6), pp.593–605.

Celis-Morales, C.A. et al., 2017. Association between active commuting and incident cardiovascular disease, cancer, and mortality: prospective cohort study. BMJ, 357.

Oliphant, T.E., 2007. Python for Scientific Computing. Computing in Science & Engineering, 9(3), pp.10–20.

Millard, L.A.C., 2017. PHESANT: a tool for performing automated phenome scans in UK Biobank. International Journal of Epidemiology, (2017), pp 1–7.

Richardson, 2012. Beautiful Soup, https://www.crummy.com/software/BeautifulSoup/

McKinney, W., 2010. Data Structures for Statistical Computing in Python. In S. Van der Walt & J. Millman, eds. Proceedings of the 9th Python in Science Conference (SciPy). pp. 51–56.

Guo, W. et al., 2015. Physical activity in relation to body size and composition in women in UK Biobank. Annals of Epidemiology, 25(6), pp.406–413.

Aykan NF., 2015. Red Meat and Colorectal Cancer. Oncology Reviews, 9(1):288.

Zhang X, Shu X-O, Xiang Y-B, et al, 2011. Cruciferous vegetable consumption is associated with a reduced risk of total and cardiovascular disease mortality. The American Journal of Clinical Nutrition, 94(1), pp. 240–246.

Crowe, F. L., Appleby, P. N., Travis, R. C., Key, T. J., 2013; Risk of hospitalization or death from ischemic heart disease among British vegetarians and nonvegetarians: results from the EPIC-Oxford cohort study, The American Journal of Clinical Nutrition, 97(3), pp. 597–603.

Bradbury, K., Tong, T., Key, T., Bradbury, K. E., Tong, T. Y. N., & Key, T. J., 2017. Dietary Intake of High-Protein Foods and Other Major Foods in Meat-Eaters, Poultry-Eaters, Fish-Eaters, Vegetarians, and Vegans in UK Biobank. Nutrients, 9(12), 1317.

Tong, T. Y., Key, T. J., Sobiecki, J. G., & Bradbury, K. E., 2018. Anthropometric and physiologic characteristics in white and British Indian vegetarians and nonvegetarians in the UK Biobank. American Journal of Clinical Nutrition, 107(6), pp. 909–920.

Hanscombe K. B., et al, 2017. ukbtools: An R package to manage and query UK Biobank data. BioRxiv 158113

